## Supplementary material for "Investigations of the Underlying Mechanisms of HIF-1*α* and CITED2 Binding to TAZ1": SI Appendix

September 2, 2019


Wen-Ting Chu[1], Xiakun Chu[2], Jin Wang[1,2*]

[1]*State Key Laboratory of Electroanalytical Chemistry, Changchun Institute of Applied Chemistry, Chinese Academy of Sciences, Changchun, Jilin, 130022, China*

[2]*Department of Chemistry & Physics, State University of New York at Stony Brook, Stony Brook, NY, 11794, USA*

### 1 Materials and Methods

#### 1.1 Initial model of simulation

NMR structures 1L8C[1] and 1R8U[2] were used for preparing the initial models of TAZ1-HIF-1$\alpha$ and TAZ1-CITED2 complexes. Full-length HIF-1$\alpha$ and CITED2 are 51 and 50 a.a. proteins (776-826 and 220-269), respectively. The initial coarse-grained $C_\alpha$ structure-based model (SBM) of TAZ1-HIF-1$\alpha$ and TAZ1-CITED2 complexes was generated using SMOG on-line toolkit[3–6]. There are 3 $Zn^{2+}$ ions linked with TAZ1 with coordination bonds, modeled by one bead with 2 positive charge (+2e) for each ion. In the present work, the weighted contact map was built with all the 20 configurations in each NMR structure. Each native contact was identified by the CSU algorithm[7]. The weighted coefficient (for intermolecular contacts and the contacts within HIF-1$\alpha$/CITED2) is the frequency of occurrence in all the configurations, similar as the method in our previous studies[8]. The potential energy function consists of both bonded and non-bonded terms. Additionally, we introduced the charge characteristics into our SBM model to study the electrostatic interactions in this system. As a result, the potential energy form used in this study is given in the following equation:

$$
\begin{aligned}
V = & \sum_{bonds} \varepsilon_r (r - r_0)^2 + \sum_{angles} \varepsilon_h \varepsilon_\theta (\theta - \theta_0)^2 \\
& + \sum_{dihedral} \varepsilon_h \varepsilon_\phi^{(n)} (1 - \cos(n \times (\phi - \phi_0))) \\
& + \sum_{contacts} \varepsilon_{ij} \left( 5 \left( \frac{\sigma_{ij}}{r_{ij}} \right)^{12} - 6 \left( \frac{\sigma_{ij}}{r_{ij}} \right)^{10} \right) \\
& + \sum_{non-contacts} \varepsilon_{NC} \left( \frac{\sigma_{NC}}{r_{ij}} \right)^{12} + \varepsilon_{DH} V_{Debye-Hückel}
\end{aligned}
\tag{S1}
$$

In Eq. S1, $\varepsilon_r = 100\varepsilon$, $\varepsilon_\theta = 20\varepsilon$, $\varepsilon_\phi^{(1)} = \varepsilon$ and $\varepsilon_\phi^{(3)} = 0.5\varepsilon$.

#### Electrostatic interactions

The electrostatic interactions are calculated by the $Debye-Hückel$ model, which can quantify the strength of charge-charge attraction and repulsion at various salt concentrations:

$$V_{Debye-Hückel} = \Gamma_{DH} \times K_{coulomb} B(\kappa) \sum_{i,j} \frac{q_i q_j \exp\left(-\kappa r_{ij}\right)}{\varepsilon r_{ij}} \tag{S2}$$

In Eq. S2, $K_{coulomb} = 4\pi\varepsilon_0 = 138.94\ kJ \cdot mol^{-1} \cdot nm \cdot e^{-2}$ is the electric conversion factor; $B(\kappa)$ is the salt-dependent coefficient; $\kappa^{-1}$ is the Debye screening length, which is directly influenced by the solvent ion strength (IS)/salt concentration $C_{salt}$ ($\kappa \approx 3.2\sqrt{C_{salt}}$); $\varepsilon$ is dielectric constant, which is set to 80 during the simulations. $\Gamma_{DH}$ is the energy scaled coefficient which aims to make the total energy balanceable. In our model, Lys and Arg have a positive point charge (+e), Asp and Glu have a negative point charge (-e). All the charges are placed on the $C_\alpha$ atoms. Under the physiological ionic strengths ($C_{salt} = 0.15M$), $\kappa$ is 1.24 $nm^{-1}$, so we set $\Gamma_{DH} = 0.535$ in our simulations, so that $V_{DH}$ for two opposite charged atoms located at a distance of 0.5 nm matches the native contact energy. When a native contact is an ionic pair (salt bridge), we rescaled its interaction strength by setting $\varepsilon_{DH} = 0.1$ so that its energetic contribution will be comparable to other native contacts[9]. More details of $Debye-Hückel$ model can be found in these papers[10–13].

#### 1.2 Parameter calibration

For angle and dihedral terms, some hinge regions were defined according to the structural flexibility of the NMR data, aiming to collect the information of conformational change during ligand binding. In this method, local interactions are weakened by decreasing the site-specific constants from the previous studies[13,14]. When variances of angle and dihedral are higher than 12.82 and 40.50 degrees, the potential energies of them are higher than 1.0 kJ/mol. Because the structure of TAZ1 is stable among the NMR structures of each complex, here we calculated the hinge regions of TAZ1 between TAZ1-HIF-1$\alpha$ and TAZ1-CITED2 to ensure the conformational flexibility of TAZ1 in the ternary complex. Then the hinge regions of HIF-1$\alpha$ or CITED2 were measured within their structures in 1L8C or 1R8U, respectively. In this model, if the angle or dihedral belong to the hinge regions, $\varepsilon_\theta$ or $\varepsilon_\phi^{(n)}$ is rescaled by setting $\varepsilon_h = 0.01$, mimicking the flexibility. Otherwise $\varepsilon_h = 1$.

For the non-local attractive term (Lennard-Jones potential) in potential energy, we divided it into five parts: intra-TAZ1, intra-HIF-1$\alpha$, intra-CITED2 terms, as well as the inter-molecular terms between TAZ1 and HIF-1$\alpha$, between TAZ1 and CITED2. These parts have different parameters of Lennard-Jones potential ($\varepsilon_{ij}$):

$$V_{non-bond}^{native} = \alpha_T V_{intra}^{TAZ1} + \alpha_H V_{intra}^{HIF-1\alpha} + \alpha_C V_{intra}^{CITED2} + \beta_H V_{inter}^{TAZ1-HIF-1\alpha} + \beta_C V_{inter}^{TAZ1-CITED2} \quad \text{(S3)}$$

As shown in Eq. S3, the $\alpha$ parameters are set for intra-molecular terms and the $\beta$ parameters are set for inter-molecular terms. The strength $\alpha_T$ was set to 1.0. Other intra-molecular parameters $\alpha_H$ and $\alpha_C$ were tuned according to the helical content of HIF-1$\alpha$ and CITED2 at unbound state. The inter-molecular parameters $\beta_H$ and $\beta_C$ were tuned according to the dissociation constant $K_d$ in experiment.

There are a little differences between the structures of TAZ1 in 1L8C and 1R8U. Therefore we kept the intra-TAZ1 native contacts with the ratio of distances $R_{ij}$ in 1L8C and 1R8U in the range of 0.8 to 1.25, and discarded the other native contacts to make the TAZ1 more "flexible". For the MD simulation with structure-based model was run with reduced units, the simulation temperature should be calibrated firstly. However, there is no melting temperature of TAZ1 reported. Because there is a critical phenomenon that the three $Zn^{2+}$ ions stabilize the structure of TAZ1, we built 4 different models of TAZ1 (TAZ1 with 3 $Zn^{2+}$, 2 $Zn^{2+}$, 1 $Zn^{2+}$, as well as TAZ1 without $Zn^{2+}$) to find out the effect of $Zn^{2+}$ on the thermodynamic stability of TAZ1. As shown in Fig. S7, at temperature about 0.99, TAZ1 with 3 $Zn^{2+}$ is stable but TAZ1 without $Zn^{2+}$ unfolds. As a result, the simulation temperature is set to 0.99, mimicking the room temperature.

In the experiment, the isolate HIF-1$\alpha$ or CITED2 was considered as "random coil"[1,2]. Here we calibrated the model to make the helical content of HIF-1$\alpha$/CITED2 below 10%. The helical content can not decline by just altering the strength of the native contact within HIF-1$\alpha$/CITED2. We then adjusted the dihedral potential of HIF-1$\alpha$ and CITED2 empirically by adding a term $V(\phi) = k_\phi cos[\phi - \delta]$, where $\delta = 297.35°$ [15]. We tuned the value of $k_\phi$ and found that when it equals to 1.0 or 0.9, the helical content of HIF-1$\alpha$/CITED2 is about 10%.

In order to achieve sufficient sampling, TAZ1-HIF-1$\alpha$ and TAZ1-CITED2 were placed in a sphere with a radius of 6 nm, leading to an effective concentration for the components of TAZ1 ($\left[C^0\right]^{Sim}$) about 1.83 mM ($\left[C^0\right]^{Sim} = \frac{1660}{V_0}$, where $V_0$ is the box volume in units of Å$^3$, 1660 is the unit transferring constant from units of molecules per Å$^3$ to units of mol/L, similar settings can be also found in previous papers[13,16–18]). Therefore,

$$K_d = \frac{[L][R]}{[LR]} = \frac{\left(P_{ub}\left[C^0\right]^{Sim}\right)\left(P_{ub}\left[C^0\right]^{Sim}\right)}{P_b\left[C^0\right]^{Sim}} = \frac{P_{ub}^2\left[C^0\right]^{Sim}}{P_b} \quad \text{(S4)}$$

where $P_{ub}$ and $P_b$ are the fractions of population of unbound states and bound states at equilibrium, respectively. From the experimental $K_d$, we can obtain the ratio of $P_{ub}/P_b$ at simulation condition (initial concentration $\left[C^0\right]^{Sim}$). Then we can obtain $\Delta G$ by applying the Boltzamann distribution $\Delta G = kT\ln(P_{ub}/P_b)$. At equilibrium of the simulations, $P_b$ is far larger than $P_{ub}$ and close to 1. As a result, when the experimentally determined $K_d$ between TAZ1 and HIF-1$\alpha$/CITED2 is 10 nM[19], the binding free energy between ligand and TAZ1 in Eq. S4 is about -6.06 kT in our model, if the effective simulation concentrations were applied. The method of binding free energy calculation is the same as that in the previous papers[15–18,20]. The strengths of inter-molecular interactions ($\beta_H$ and $\beta_C$) were tuned by performing a series of REMD simulations on TAZ1-HIF-1$\alpha$ and TAZ1-CITED2 complexes, respectively. As shown in Fig. S8, both $\beta_H$ and $\beta_C$ are set to be 1.1 and 0.95 in our model.

In the ternary system, we can obtain the simulated $K_d$ by using the free energy difference between bound state (HB or CB) and unbound state (UB).

#### 1.3 MD simulation

All simulations were performed with Gromacs 4.5.5[21]. The coarse grained molecular dynamics simulations (CGMD) used Langevin equation with constant friction coefficient $\gamma = 1.0$. The cutoff length for non-bonded interactions was set to 3.0 nm. The MD time step was set to 0.5 fs and the trajectories were saved every 2 ps. To enhance the sampling of binding events, a strong harmonic potential was added if the distance between the center of mass of TAZ1 and HIF-1$\alpha$, TAZ1 and CITED2 is greater than 6 nm[22].

For thermodynamic simulations (binding and unbinding for multiple times), REMD simulations and long-time MD simulations were performed to overcome the energy barriers between bound and unbound states. We define that a native contact is formed if the C$\alpha$-C$\alpha$ distance between any given native atom pair is within 1.2 times of its native distance. Then the profiles of free energy curve or surface can be obtained by using WHAM algorithm[23,24]. The REMD simulations have been tested to be converged that the fraction of all the native contacts in the ternary system (Q) becomes equilibrated and stable after about 150 ns of each replica (see Fig. S9).

For kinetic simulations, 200 individual MD runs started with varying configurations and velocities were performed on different processes respectively: direct binding (both unbound state to TAZ1-HIF-1$\alpha$ or TAZ1-CITED2 state) and replacement (CITED2 binding to TAZ1 by replacing HIF-1$\alpha$ and HIF-1$\alpha$ binding to TAZ1 by replacing HIF-1$\alpha$).

#### 1.4 $\phi$ value of binding

The calculation of $\phi$ value is referred to the previous simulation papers[4,25]. The $\phi_{ij}$ for each inter-molecular native contact pair between residue $i$ and $j$ was computed from the probability of formation, $P_{ij}$:

$$\phi_{ij} = \frac{\Delta\Delta F^{TS-U}}{\Delta\Delta F^{B-U}} \approx \frac{P_{ij}^{TS} - P_{ij}^{U}}{P_{ij}^{B} - P_{ij}^{U}} \tag{S5}$$

where $\Delta\Delta F$ is the free energy difference between the wild-type and mutated protein, $P_{ij}$ is the probability of formation of contact between $i$ and $j$. Here, U, TS, B correspond to the unbound state, transition state ($Q_{inter} \sim 0.03 - 0.1$), and bound state, respectively. Then, $\phi_i$ value of residue $i$ can be calculated from the average of $\phi_{ij}$ that are involved with residue $i$.

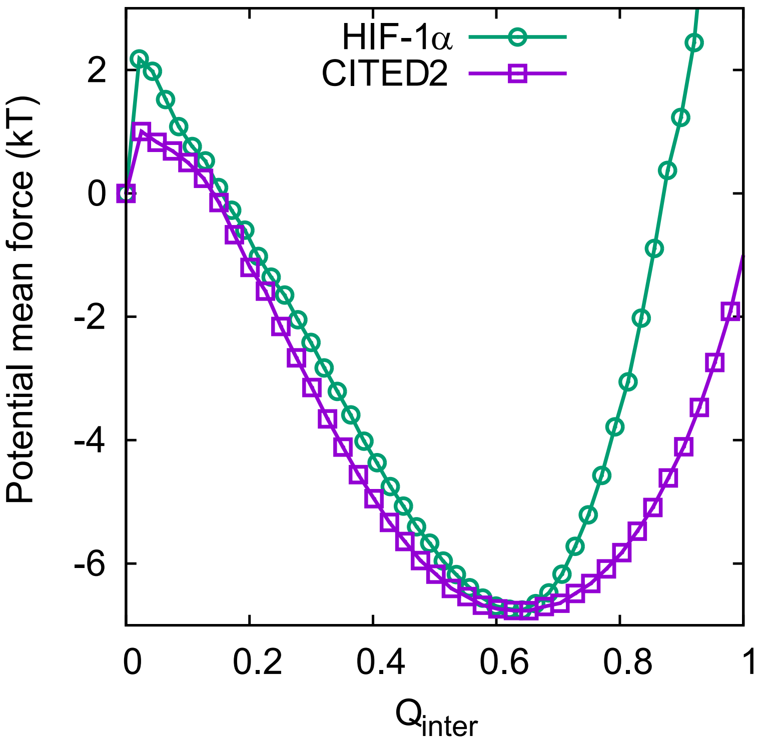


**Fig. S1** The free energy curves as a function of the fraction of inter-molecular native contacts ($Q_{inter}$) of TAZ1-HIF-1$\alpha$ (green line) and TAZ1-CITED2 (magenta line). The free energy unit is kT (k is Boltzmann constant).

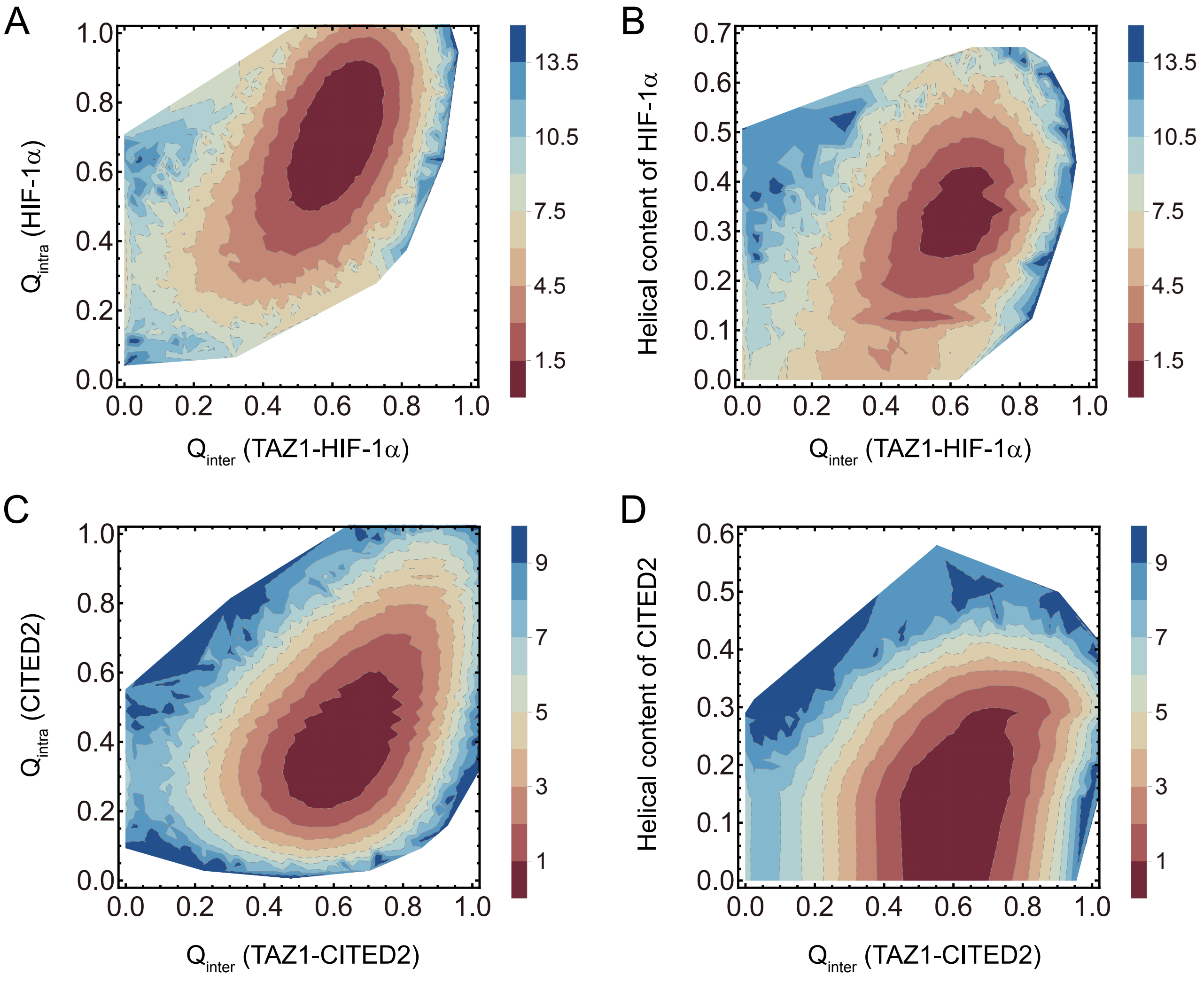


**Fig. S2** The free energy surfaces projected on (A) both the fraction of inter-molecular native contacts ($Q_{inter}$) of TAZ1-HIF-1$\alpha$ and the fraction of intra-molecular native contacts ($Q_{intra}$) of HIF-1$\alpha$; (B) both $Q_{inter}$ of TAZ1-HIF-1$\alpha$ and the helical content of HIF-1$\alpha$; (C) both $Q_{inter}$ of TAZ1-CITED2 and $Q_{intra}$ of CITED2; (D) both $Q_{inter}$ of TAZ1-CITED2 and the helical content of CITED2. The free energy unit is kT (k is Boltzmann constant).

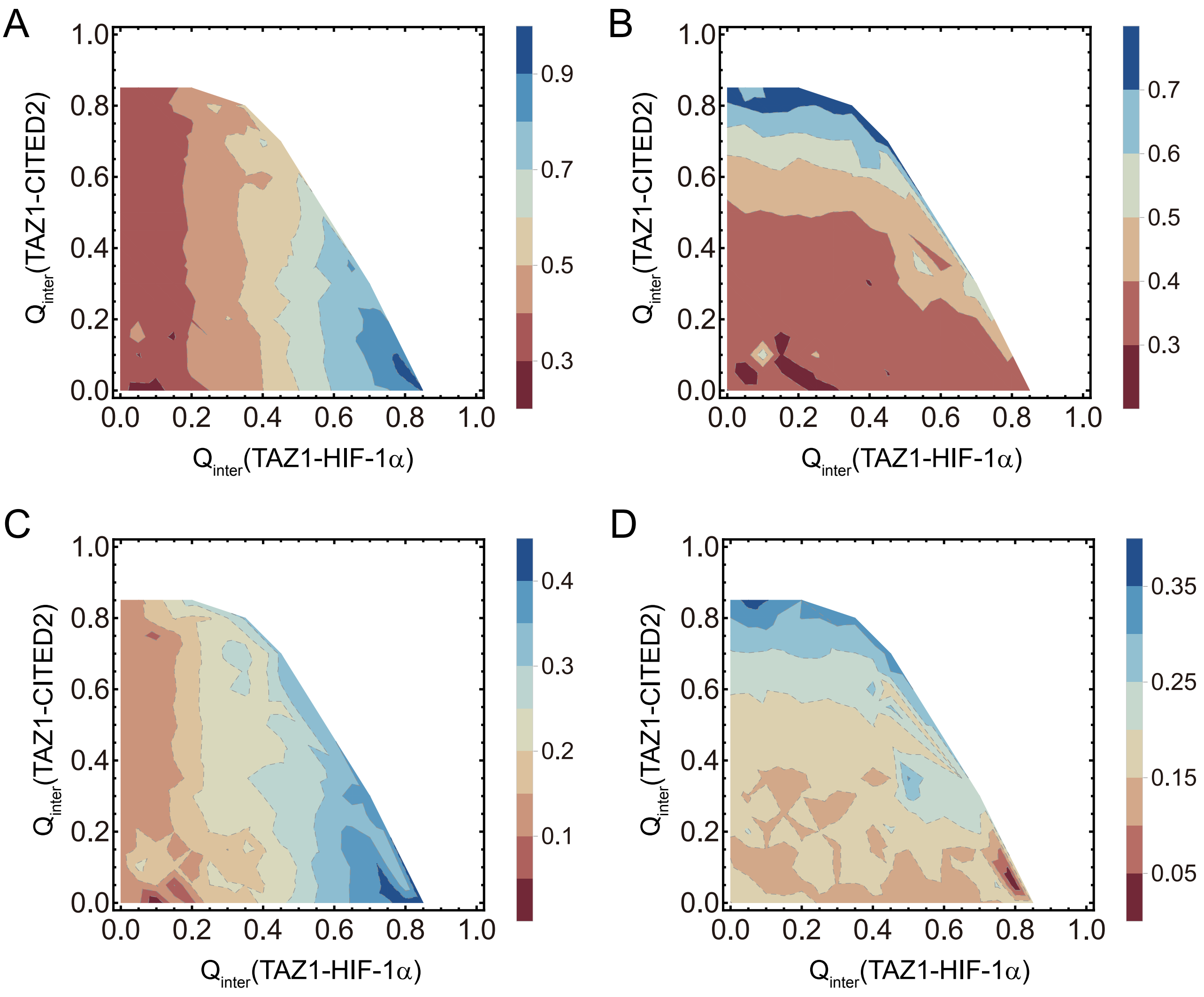


**Fig. S3** Mean $Q_{intra}$ of HIF-1$\alpha$ (A) and CITED2 (B) as well as mean helical content of HIF-1$\alpha$ (C) and CITED2 (D) as a function of $Q_{inter}$ (TAZ1-HIF-1$\alpha$) and $Q_{inter}$ (TAZ1-CITED2).

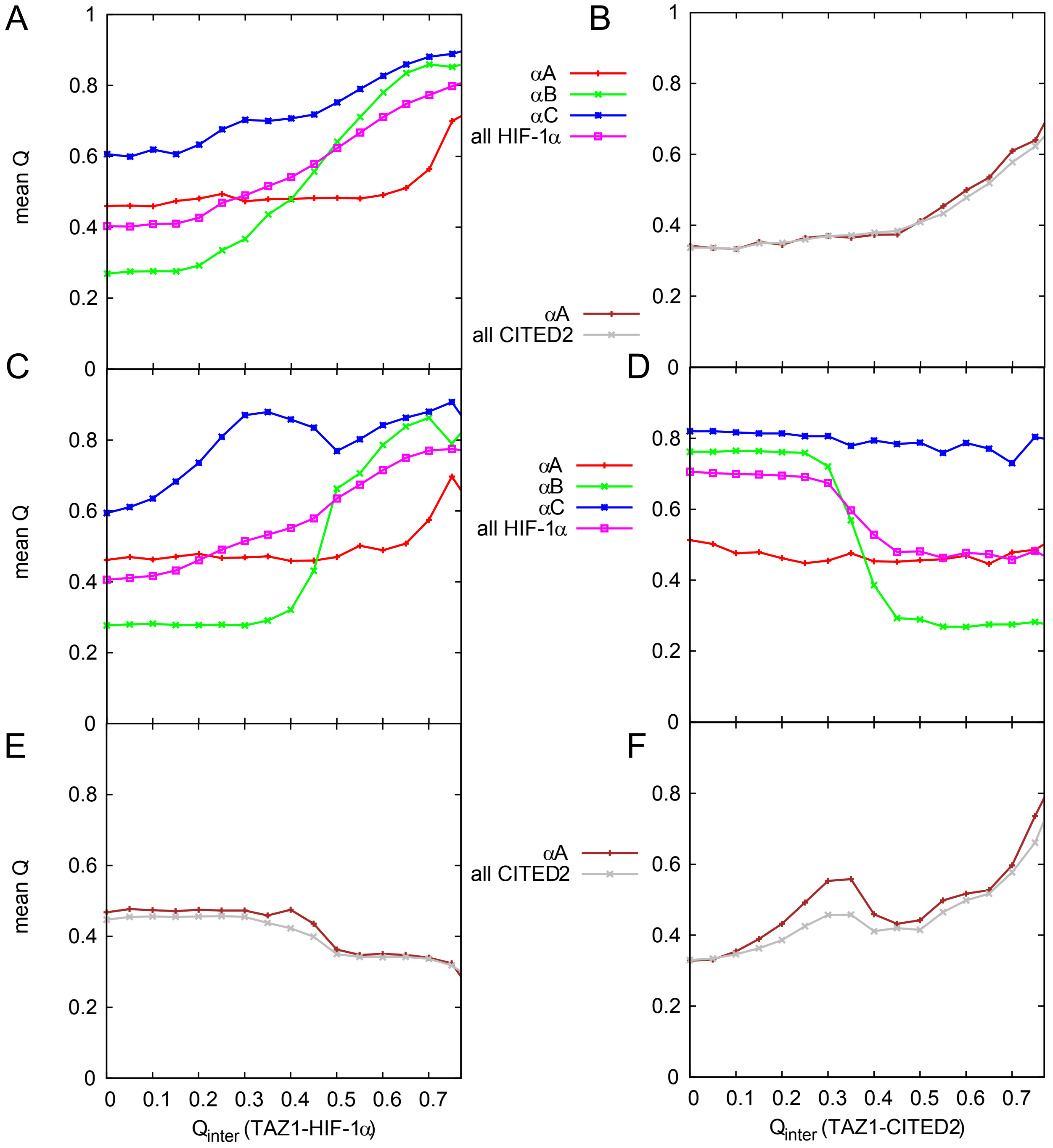


**Fig. S4** The mean $Q_{intra}$ within different parts of ligand (HIF-1$\alpha$ or CITED2) as a function of binding in the UH (A), UC (B), CIH (C and E), and HIC (D and F) pathways. Panels A, C, and D show the $Q_{intra}$ curves of HIF-1$\alpha$; Panels B, E, and F show the $Q_{intra}$ curves of CITED2.

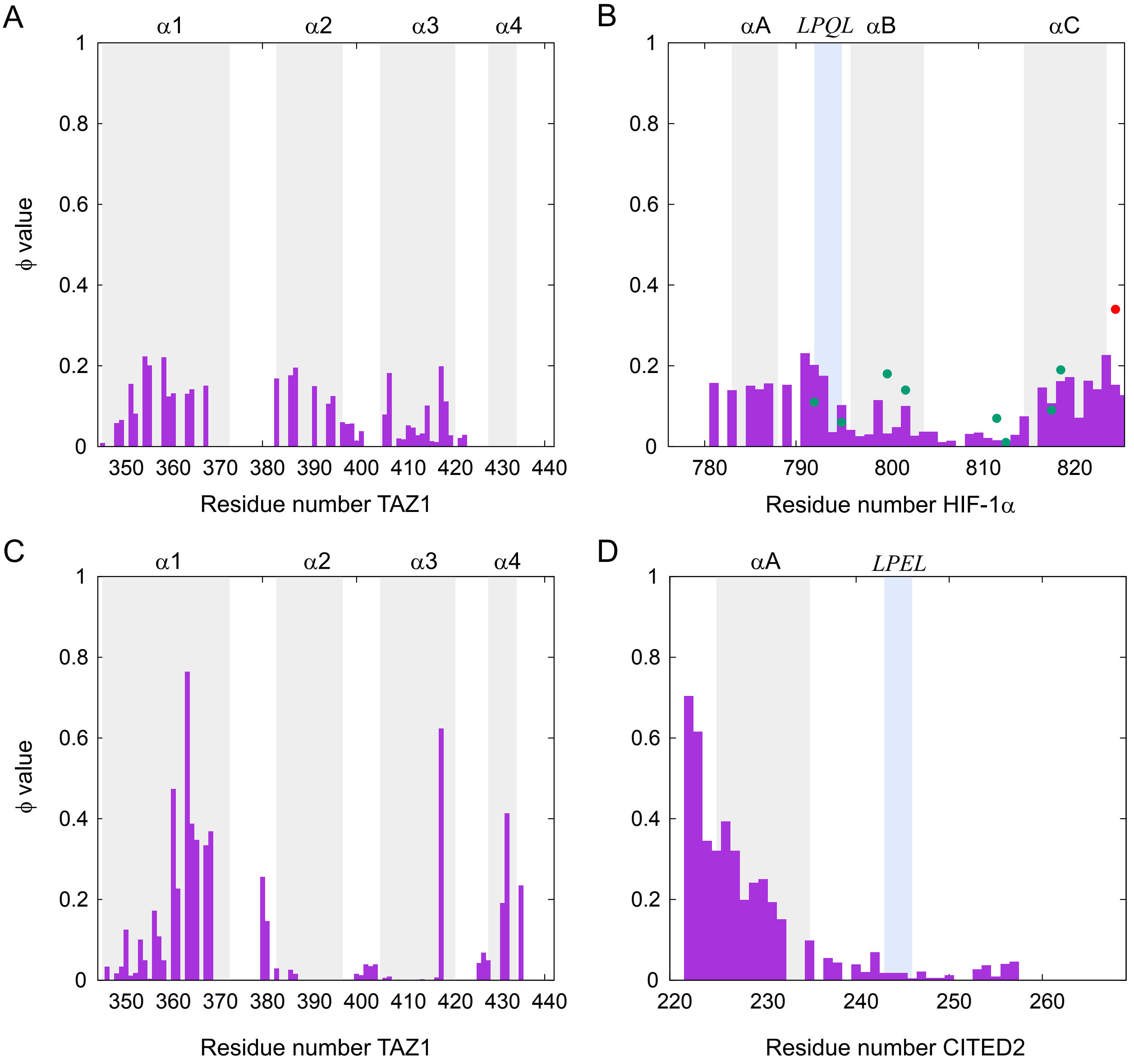


**Fig. S5** The $\phi$ values of TAZ1 (residue 345 to 439, as well as 3 $Zn^{2+}$) in UH (A) and UC (C) pathways as well as the $\phi$ values of HIF-1$\alpha$ (residue 776 to 826) in UH pathway (B) and CITED2 (residue 220 to 269) in UC pathway (D). The experimental $\phi$ values (in ref[26]) are shown in dots. The secondary structures as well as the LPQL/LPEL motif are labeled.

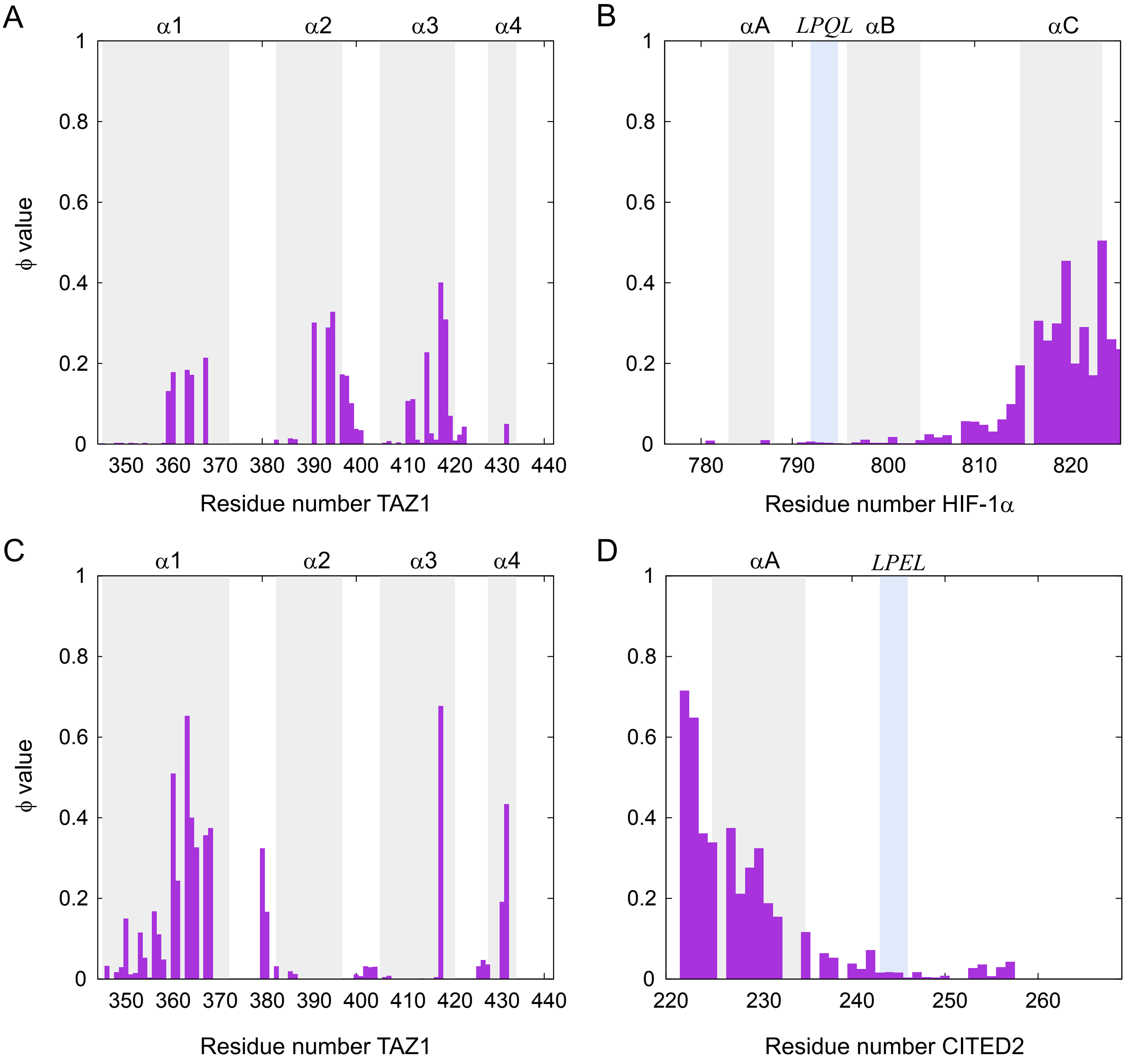


**Fig. S6** The $\phi$ values of TAZ1 (residue 345 to 439, as well as 3 $Zn^{2+}$) in CIH (A) and HIC (C) pathways as well as the $\phi$ values of HIF-1$\alpha$ (residue 776 to 826) in CIH pathway (B) and CITED2 (residue 220 to 269) in HIC pathway (D). The secondary structures as well as the LPQL/LPEL motif are labeled.

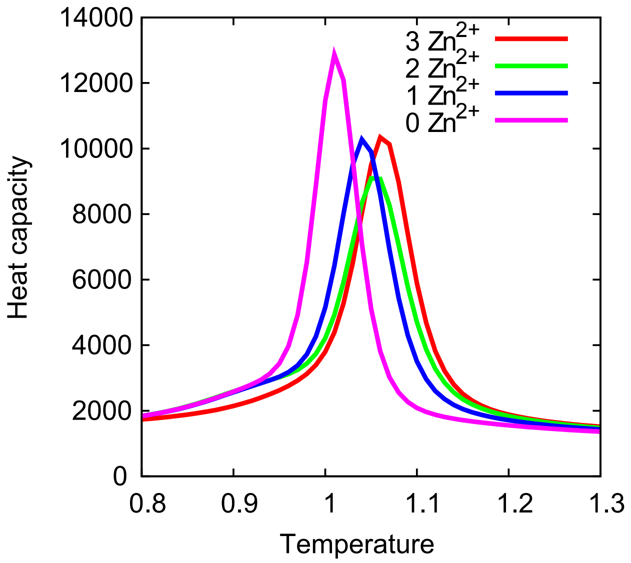


**Fig. S7** The heat capacity curves of TAZ1 with 3 $Zn^{2+}$ (red), 2 $Zn^{2+}$ (green), 1 $Zn^{2+}$ (blue), as well as TAZ1 without $Zn^{2+}$ (magenta).

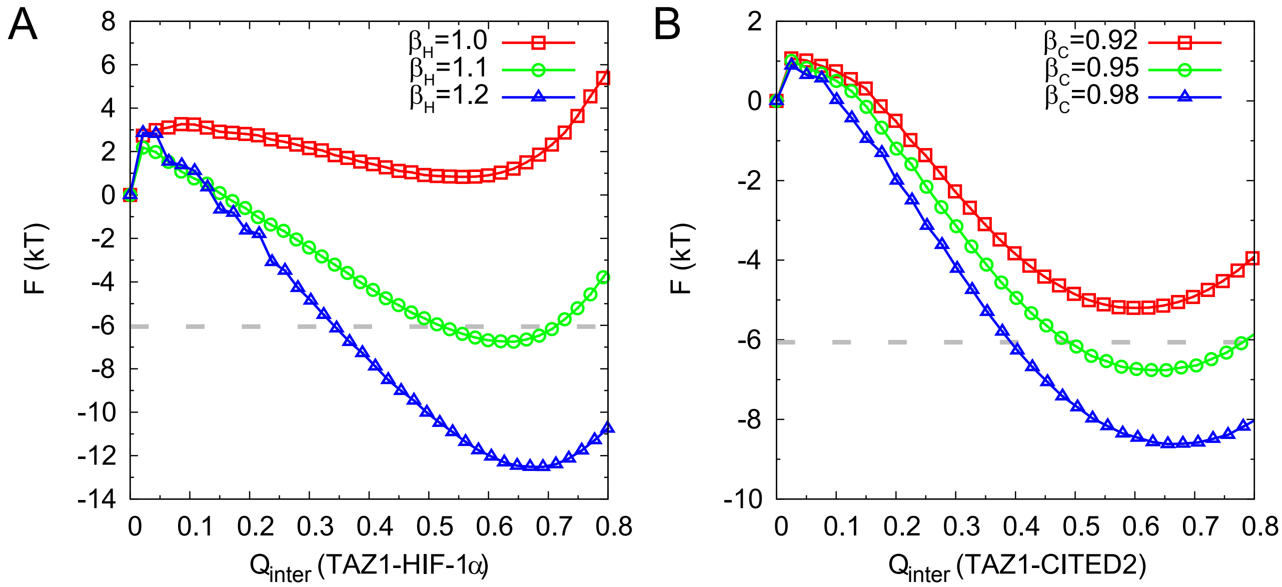


**Fig. S8** Free energy as a function of HIF-1$\alpha$ binding (A) and CITED2 binding (B) with different parameters.

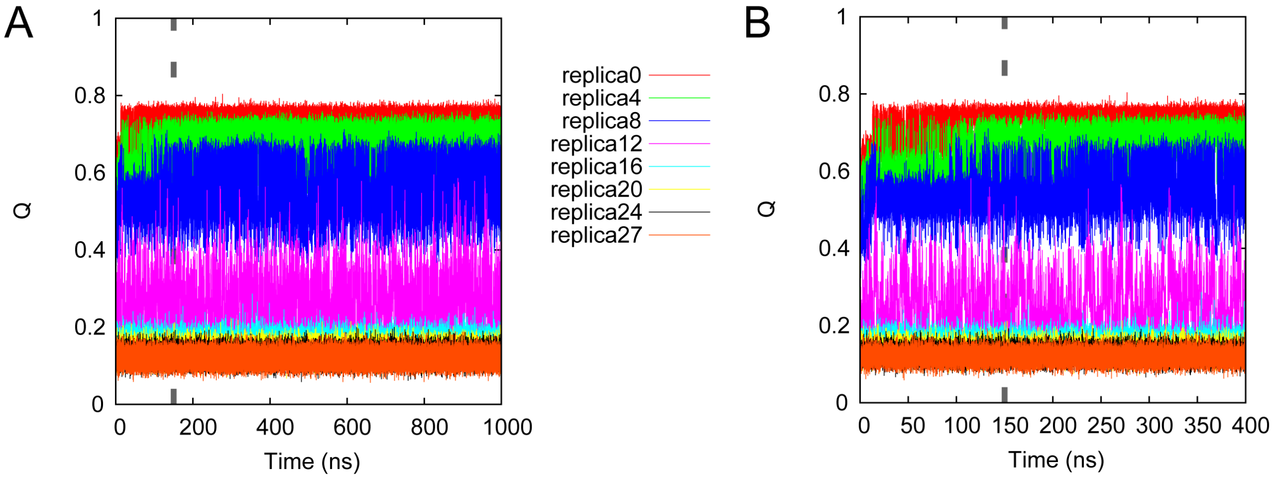


**Fig. S9** The fraction of all the native contacts in the ternary system as a function of simulation time of the REMD run (total 1 $\mu$s per replica). The native contacts include the intra-molecular (within TAZ1, HIF-1$\alpha$, and CITED2) and the inter-molecular (between TAZ1 and HIF-1$\alpha$, between TAZ1 and CITED2) contacts. The right panel (B) is the first 400 ns.
